# Prolonged survival of rabid mice after treatment with Poly-ICLC Hiltonol®

**DOI:** 10.1101/2023.09.18.558218

**Authors:** P.E. Brandão, W.C. Agostinho, O.L. Caballero, A.M. Salazar, C.O.M.S. Gomes

## Abstract

Despite its 100% lethality and approximately 59,000 human deaths a year, rabies is still in want of an effective treatment. A host of trials has been described aiming at impairing the life cycle of *Lyssavirus rabies* (RABV), the chief worldwide lyssavirus causing rabies, but with limited success. In this study, mice intranasally inoculated with the CVS strain of RABV were serially treated with 50µg/day of the Poly ICLC Hiltonol® jn MEM showed significantly higher survival time and disease incubation period as compared to control mice that received MEM only. These very preliminary results indicate that Hiltonol® might be an effective drug against rabies that merits wider *in vivo* investigation.

## 1. Introduction

Human rabies is currently present in more than 150 countries in which approximately 59,000 human deaths occur every year due to the disease (https://www.who.int/health-topics/rabies#tab=tab_1). The clinical manifestation of rabies in humans, after an incubation period that ranges from some weeks to one year, starts with the prodromal phase, during which neuropathic pain that might spread to large portions of the muscular system and pruritus at the site of virus entry are amongst the early signs. The acute neurological phase exhibits autonomic dysfunction, hyperesthesia, and fluctuations in consciousness. Manifest, aggressiveness and hallucinations characterize the furious form of the disease, while a progressive muscle weakness is seen in cases of the paralytic form of the disease (Hemachudha and Hemachudha, 2021).

Despite in-depth knowledge on the causative agent (*Lyssavirus rabies* RABV, *Mononegavirales*: *Rhabdoviridae*: *Lyssavirus*) and the centuries of trial and (mostly) error approaches, no efficient specific treatment is available for those who have not undergone post-exposure prophylaxis based on inactivated vaccines and hyperimmune sera. A valid treatment for rabies must be effective in patients with clinical manifestations of the disease and should not be mistaken with post or pre-exposure preventive treatments based on vaccines that, although highly effective, are of no use when the RABV is already present in the patient’s CNS.

RABV has evolved several mechanisms to evade or manipulate the host’s innate immune response, such as the inhibition of interferon production by suppressing activation of pattern recognition receptors or inhibiting signalling pathways (Lafon, 2011).

It has been shown that recombinant human interferon alpha prolongs survival of rabies infected mice (Mehta et al., 2015). Furthermore, mice peripherally (*i*.*e*., not into the central nervous system) treated with two injections of poly I:C (3- and 15-hours post-infection) exhibit an increase in IFN levels and are significantly protected (Harmon et al., 1975). This apparent efficacy, however, may have been due to the short period between inoculation and treatment.

A modified form of Poly I:C with carboxymethylcellulose and polylysine as stabilizers (Poly ICLC), has been shown to be more effective in inducing IFN in primates as compared to Poly I:C alone both with (Baer et al., 1979) and without RABV inoculation (Levy et al., 1975). It should be noted that Baer et al used the intramuscular route for RABV inoculation, as this could result in abortive infections.

Hiltonol® is an experimental Poly ICLC viral mimic and broad activator of innate and adaptive immunity with interferon-inducing properties and has been used as an adjuvant in the setting of peptide-based cancer vaccines and alone for in-situ tumour autovaccination (Sultan et al., 2020) and as an antiviral against SARS-CoV-1 and Ebolavirus (Kumaki et al., 2017; Kende et al., 2019), among other viruses.

The aim of this preprint is to report the effect of Hiltonol® on the survival rates and incubation periods in mice with central nervous system infection.

## 2. Materials and Methods

### 2.1 Inoculation of mice with RABV and treatment protocol

Eighteen 60-day-old mice were inoculated via the intranasal route with 10µL per nostril of the CVS strain of RABV (at 10^5.3^ TCID_50_/mL) at Day 0 and then separated in two groups, *i*.*e*., Treated and Control groups. All procedures with mice were carried under Isoflurane anaesthesia. Mice were kept with feed and water ad libitum and under a 12h light regimen and the experiments as per the norms approved by National Council for Control of Animal Experimentation (Concea), Brazil.

Form days three to 14, mice in the Treated group received 2mL Hiltonol® 25µg/mL (50µg/mouse) in MEM (Minimum Essential Medium) intraperitoneally *q*.*d*., while the Control group was injected *q*.*d*. MEM only, both warmed to room temperature prior to injection. MEM was selected as a vehicle to provide parenteral hydration and feeding as the mice are unable to drink and feed due to rabies.

Mice were observed daily for signs of two stages of rabies: stage S-1: ruffled hair and weight loss; stage S-2: ataxia, paralysis, and tremors. Dead mice were frozen at -20 ºC to preserve the central nervous system for the subsequent analysis.

### 2.2 Direct fluorescent antibody test (dFAT) and real time PCR

The central nervous systems of mice were collected after opening the skulls and assayed for RABV antigens using an anti-RABV nucleocapsid fluorescein isothiocyanate conjugate on acetone-fixed impressions as described by Dean et al. (1996).

Total RNA was extracted from mice brains suspensions using the Mini Kit RNA Purelink and (Life Technologies, Carlsbad, USA) and the RABV genome copy numbers assessed with a SYBR green-qPCR targeting RABV N gene with Beta-actin TaqMan Probe qPCR as a reference as described by Wakeley et al. (2005).

### 2.3 Statistical analyses

The number of days from dpi 0 to the first sign of rabies (incubation period) and to the first death (survival) in the control and treated groups were compared using Survival analysis and the Logrank and Wilcoxon tests using a critical p value of 0.05.

## 3. Results

### 3.1 Rabies signs and mortality

No signs of rabies were detected up to days 7 and 9 post-inoculation (dpi) in the Control and Treated groups, respectively. Also, no mortality was detected up to dpi 10 and 11 in the Control and Treated groups, respectively (Table 1 and Fig. 1).

**Table 1.**
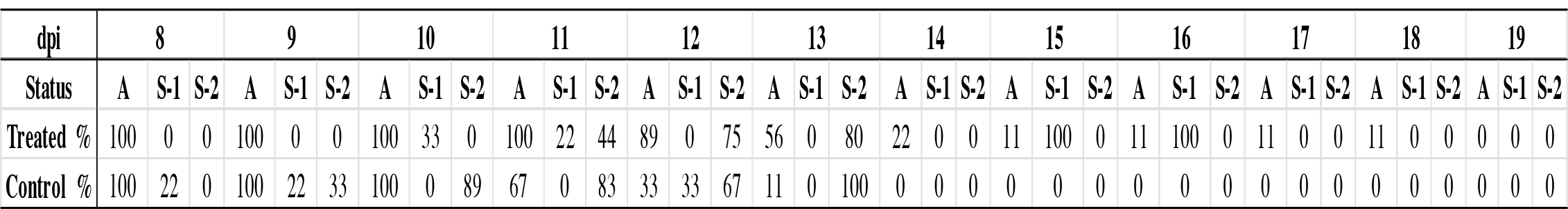
Frequency of mice inoculated via intranasal with the CVS strain of RABV and treated with 2mL Hiltonol® (25µg/mL in MEM) ou Control (2 mL MEM) at 3 days post inoculation (dpi) alive (A) or with rabies signs at stage 1 (S-1: ruffled hair and weight loss) or stage 2 (S-2: ataxia, paralysis, and tremors); nine mice were used per group. Groups were treated from dpi 3 to 14; the last surviving mouse form the Treated group was also treated at dpi 17 and 18. As no signs or mortality were detected from dpi 0 to 7, these are not shown.

**Figure 1.**
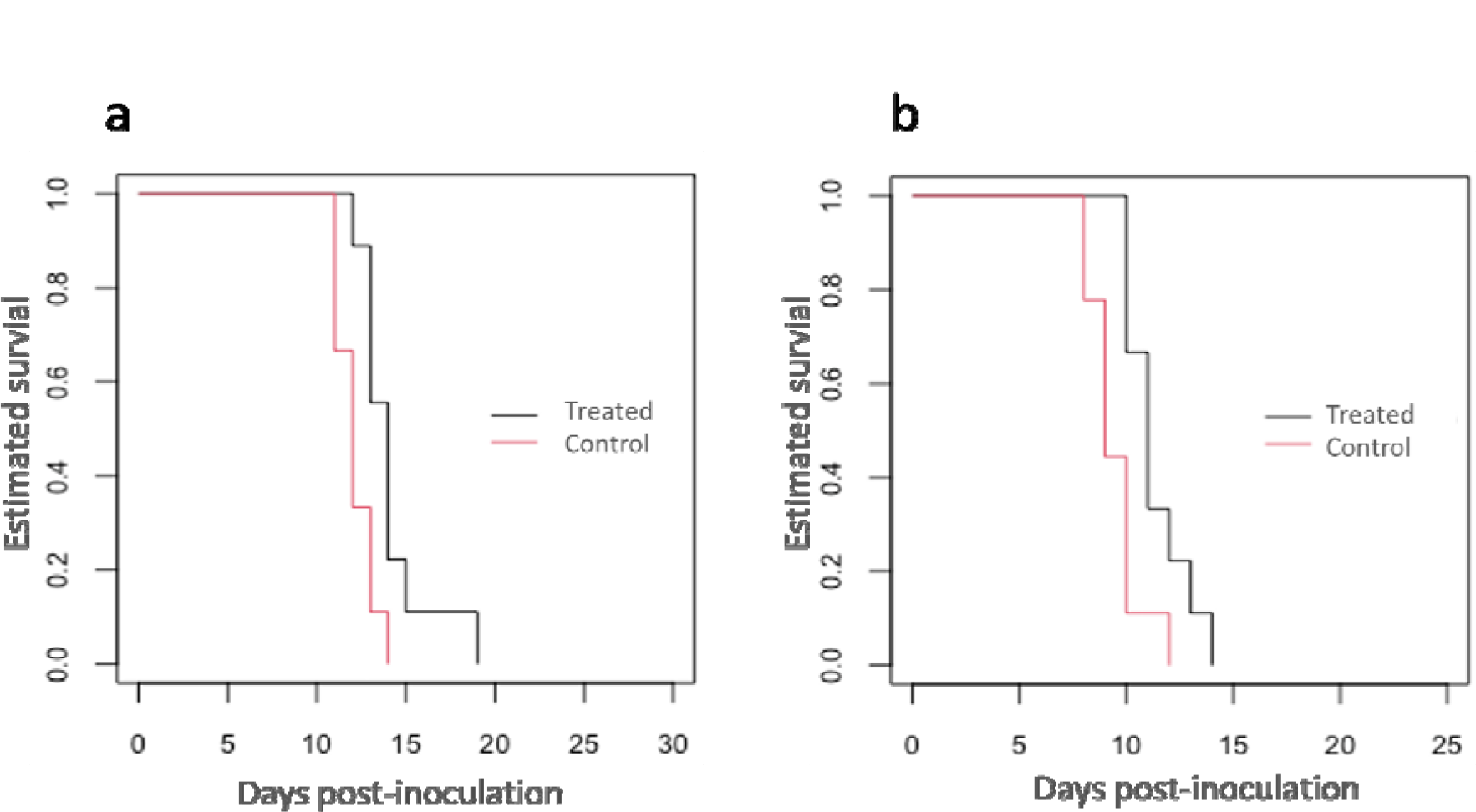
Survival analysis of mice inoculated via the intranasal route with the CVS strain of RABV and treated with 2mL Hiltonol® (25µg/mL in MEM) or 2 mL MEM at three days post inoculation (dpi) for a. deaths and b. mice with any signs of rabies; nine mice were used per group. Groups were treated (with Hiltonol or MEM) from dpi 3 to 14 and the last surviving mouse form the Treated group was also treated with 2mL Hiltonol® (25µg/mL in MEM) at dpi 17 and 18.

In the Control group, two mice (22%) showed S-1 signs at dpi 8 and two days later, eight (89%) were in the S-2 stage; the last control mouse died at the S-2 stage at dpi 14. On the other hand, signs of the S-1 level were first detected only in 3 (33%) in the Treated group at dpi 10 and the first death at 12 dpi (Table 1 and Fig. 1).

At dpi 15, only a mouse of the Treated group at the S-2 stage remained alive from both groups and was still in the same condition at dpi 17. On this and the next day it was given another round of Hiltonol® treatment. No improvement was observed and the mouse died at dpi 19 (Table 1).

Survival analysis showed that mice in the treated group had statistically significant higher incubation (time to first signs) and survival times (Logrank/Wilcoxon p-values =0.010/ 0.006 and 0.010/0.008, respectively).

### 3.2 Direct immunofluorescence and RABV qPCR

All mice were positive for RABV antigens after dFAT, with no detectable differences in distribution of fluorescent foci.

Genome copy numbers in ranged from 8 to 8.9 and 7.9 to 8.7 log/µL RNA in the control and treated groups, respectively (Figure 2). In the last surviving mouse of the Treated group that received two extra Hiltonol® injections at dpis 17 and 18, the genome copy number was 7.9 log/µL RNA.

**Figure 2.**
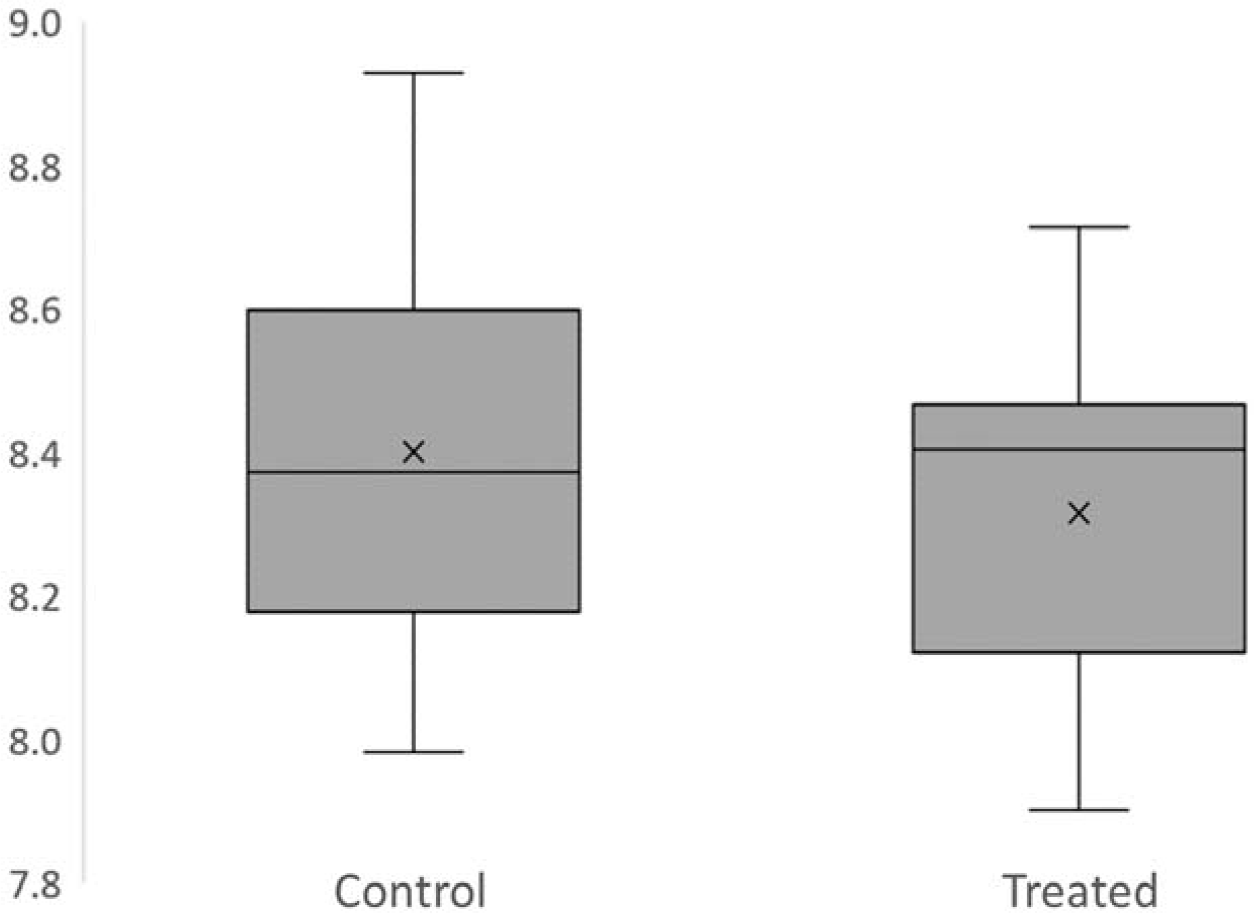
RABV genome copy numbers (log/µL RNA) in the central nervous system of mice (at time of death) inoculated via intranasal with the CVS strain of RABV and treated with 2mL Hiltonol® (25µg/mL in MEM) ou Control (2 mL MEM) at 3 days post inoculation (dpi).

## 4. Discussion

In this investigation, mice intranasally inoculated with the CVS strain of RAB and treated intraperitoneally with Hiltonol® showed a significantly higher survival time after the onset of rabies symptoms as well as a longer period between inoculation and the onset of symptoms (*i*.*e*., incubation period). Nevertheless, all mice died at dpi 19 due to rabies. In considering a treatment for rabies the elongation of the survival of a patient is of high interest as many a different parallel antivirals and supportive therapies might be administered with a view to curing the patient.

Repurposed drugs have been one of the main techniques used to find a potential antiviral drug against rabies. Ribavirin, an antiviral used in respiratory syncytial virus (RSV) infection that inhibits the production of viral messenger RNA, has been investigated and has shown *in vitro* inhibition of rabies infection. In mice, however, ribavirin shifts the immune response to an inflammatory Th1-type response, while resistance to rabies requires a balanced Th1/Th2 immune response (Appolinario and Jackson, 2014). This suggests that ribavirin might suppress antibody production which aids the clearance of rabies from the patient to some degree.

Favipravir is an RNA virus inhibitor that has also been assessed as a repurposed drug against rabies. Although it has an interesting mechanism of action and diminished signs of rabies in mice, it had no effect in the CNS, which is of the utmost importance, considering that is where rabies infection causes most harm (Du Pont et al., 2019).

In a previous experiment (unpublished data), we have found that mice inoculated with 10ten CVS strain of RAB at the same dose used herein (10^5.3^ TCID_50_/mL) showed log RABV genome copy numbers/µl of 3.95, 5.98, 7.56, 8.75, 8.56, 8.85, 8.87, 8.93 at dpis 3, 4, 5, 6, 7, 8, 9 and10, respectively, showing that RABV was already in then central nervous system when mice were first treated at dpi 3 and placing the copy numbers of both Treated and Control mice in the expected range at time of death.

Therefore, a balance of the immune system regulation could be achieved with the use of Hiltonol® a synthetic double-stranded RNA more RNase-resistant analogue stabilized with poly-L-lysine (poly ICLC) shown to mediate robust adjuvant effects via induction of type I IFNs via recognition by the cytosolic RNA helicase MDA-5 and by endosomal TLR and trigger the activation of TRAF3 and TRADD to activate TBK1 and drive type I IFN expression (Longhi et al., 2009).

One of the limitations of this study is that, as it was not possible to sample the mice daily, the evolution of the RABV genome copy numbers along with the clinical course in each animal could not be assessed and thus the precise effect of the treatment on virus replication might have been lost. This could be improved in future investigations using oral swabs. In addition, IFN was not measured in mice, and this will be of significant importance to potentially account for the successful outcome of the use of Hiltonol®. A suggestion for the improvement of the survival rate is an increase of the Hiltonol® dosage used herein (50µg/mouse) to a higher level coupled with different titres and strains of RABV.

In conclusion, Hiltonol® can be recommended for a wider *in vivo* investigation as a potential treatment of rabies.

## 5. Acknowledgements

The authors are grateful to FAPESP (São Paulo Research Foundation, grant # 2022/07115-7) and CNPq (Brazilian National Council for Scientific and Technological Development, grant # 302503 2021-8) for the financial support.

## 6. Conflict of interest

Andres M. Salazar isa n officer and owns stock in Oncovir.

## References

Appolinario CM, Jackson AC. Antiviral therapy for human rabies. Antivir Ther. 2015;20(1):1–10. doi: 10.3851/IMP2851.

Baer GM, Moore SA, Shaddock JH, Levy HB. An effective rabies treatment in exposed monkeys: a single dose of interferon inducer and vaccine. Bull World Health Organ. 1979;57(5):807–13.

Dean DJ, Abelseth MK, Atanasiu P. The fluorescent antibody test In: Koprovsky H. The laboratory techniques in rabies. Geneva. 1996. 476p.

Du Pont V, Plemper RK, Schnell MJ. Status of antiviral therapeutics against rabies virus and related emerging lyssaviruses. Curr Opin Virol. 2019 Apr;35:1–13. doi: 10.1016/j.coviro.2018.12.009.

Harmon MW, Janis B. Therapy of murine rabies after exposure: efficacy of polyriboinosinicpolyribocytidylic acid alone and in combination with three rabies vaccines. J Infect Dis. 1975 Sep;132(3):241–9. doi: 10.1093/infdis/132.3.241.

Hemachudha P, Hemachudha T. Rabies: Presentation, case management and therapy. J Neurol Sci. 2021 May 15;424:117413. doi: 10.1016/j.jns.2021.117413.

Kende M, Paragas J, Salazar AM. The efficacy of poly-ICLC against Ebola-Zaire virus (EBOV) infection in mice and cynomolgus monkeys. Antiviral Res. 2019 Mar;163:179–184. doi: 10.1016/j.antiviral.2018.12.020.

Kumaki Y, Salazar AM, Wandersee MK, Barnard DL. Prophylactic and therapeutic intranasal administration with an immunomodulator, Hiltonol® (Poly IC:LC), in a lethal SARS-CoV-infected BALB/c mouse model. Antiviral Res. 2017 Mar;139:1–12. doi: 10.1016/j.antiviral.2016.12.007.

Lafon M. Evasive strategies in rabies virus infection. Adv Virus Res. 2011; 79:33–53. doi: 10.1016/B978-0-12-387040-7.00003-2.

Longhi MP, Trumpfheller C, Idoyaga J, Caskey M, Matos I, Kluger C, Salazar AM, Colonna M, Steinman RM. Dendritic cells require a systemic type I interferon response to mature and induce CD4+ Th1 immunity with poly IC as adjuvant. J Exp Med. 6;206(7):1589–602, 2009. doi: 10.1084/jem.20090247.

Levy HB, Baer G, Baron S, Buckler CE, Gibbs CJ, Iadarola MJ, London WT, Rice J. A modified polyriboinosinic-polyribocytidylic acid complex that induces interferon in primates. J Infect Dis. 1975 Oct;132(4):434–9. doi: 10.1093/infdis/132.4.434.

Mehta S, Roy S, Mukherjee S, Yadav N, Patel N, Chowdhary A. Exogenous interferon prolongs survival of rabies infected mice. Virus disease. 2015 Sep;26(3):163–9. doi: 10.1007/s13337-015-0269-5.

Sultan H, Salazar AM, Celis E. Poly-ICLC, a multi-functional immune modulator for treating cancer. Semin Immunol. 2020 Jun;49:101414. doi: 10.1016/j.smim.2020.101414.

Wakeley PR, Johnson N, McElhinney LM, Marston D, Sawyer J, Fooks AR. Development of a real-time, TaqMan reverse transcription-PCR assay for detection and differentiation of lyssavirus genotypes 1, 5, and 6. J Clin Microbiol. 2005 Jun;43(6):2786–92. doi: 10.1128/JCM.43.6.2786-2792.2005.

